# Sex and age differences in cognitive bias and neural activation in response to cognitive bias

**DOI:** 10.1101/2022.02.02.478831

**Authors:** Travis E. Hodges, Grace Y. Lee, Sophia H. Noh, Liisa A.M. Galea

**Author notes:** Corresponding Author: Dr. Liisa Galea, Djavad Mowafaghian Centre for Brain Health, University of British Columbia, 2215 Wesbrook Mall, Vancouver, BC, Canada, V6T 1Z3.

## Abstract

Cognitive symptoms of depression, including negative cognitive bias, are more severe in women than in men. Current treatments to reduce negative cognitive bias are not effective and sex differences in the neural activity underlying cognitive bias may play a role. Here we examined sex and age differences in cognitive bias and functional connectivity in a novel paradigm. Male and female rats underwent an 18-day cognitive bias procedure, in which they learned to discriminate between two contexts (shock paired context A, no-shock paired context B), during either adolescence (postnatal day (PD 40)), young adulthood (PD 100), or middle-age (PD 210). Cognitive bias was measured as freezing behaviour in response to an ambiguous context (context C), with freezing levels akin to the shock paired context coded as negative bias. All animals learned to discriminate between the two contexts, regardless of sex or age. However, adults (young adults, middle-aged) displayed a greater negative cognitive bias compared to adolescents, and middle-aged males had a greater negative cognitive bias than middle-aged females. Females had greater neural activation of the nucleus accumbens, amygdala, and hippocampal regions to the ambiguous context compared to males, and young rats (adolescent, young adults) had greater neural activation in these regions compared to middle-aged rats. Functional connectivity between regions involved in cognitive bias differed by age and sex, and only adult males had negative correlations between the frontal regions and hippocampal regions. These findings highlight the importance of examining age and sex when investigating the underpinnings of negative cognitive bias and lay the groundwork for determining what age- and sex-specific regions to target in future cognitive bias studies.

**Highlights:**

- Middle-aged males had a greater negative cognitive bias than middle-aged females
- Adult rats displayed a greater negative cognitive bias compared to adolescents
- Greater neural activity in females than males in limbic and reward regions
- Greater role of the frontal cortex activation in the cognitive bias of adults
- Functional connectivity in response to cognitive bias differed by age and sex

## Introduction

Major depressive disorder (MDD) is the leading cause of disability and affects more than 320 million people worldwide (World Health Organization (WHO), 2017). Females are more likely to develop depression and have more severe symptoms of MDD compared to males (Bogren et al., 2018; Kornstein et al., 2002; Labaka et al., 2018; Sloan and Kornstein, 2003). Sex differences in MDD rates vary with age, becoming more prominent after puberty and reducing in middle age (Gutiérrez-Lobos et al., 2002). Despite these prominent sex differences, MDD studies in the neurobiology of MDD rarely considers sex as a discovery variable (reviewed in Eid et al., 2019).

Cognitive symptoms of MDD such as negative cognitive bias (the perception of ambiguous stimuli as negative) and rumination (repetitive negative thinking) are more common in females than in males (Kornstein et al., 2000; Mansour et al., 2006; Nolen-Hoeksema et al., 1999; Silverstein, 1999). Negative cognitive bias is involved in the maintenance, onset, and severity of MDD across the lifespan (Bar-Haim et al., 2007; Lau and Waters, 2017; Lee et al., 2016; Orchard and Reynolds, 2018; Platt et al., 2015). Cognitive symptoms of MDD are associated with increased relapse rates and meta-analyses find that they persist in individuals in remission from MDD (Bora et al., 2013; Bortolato et al., 2016; Hallion and Ruscio, 2011; Hasselbalch et al., 2011; Hilimire et al., 2015; LeMoult et al., 2018; Micco et al., 2014; Phillips et al., 2010). Furthermore, cognitive symptoms of MDD, such as negative cognitive bias, can predict the onset of MDD in healthy individuals with an increased risk for MDD as well as future depressive episodes in individuals with MDD, and these effects are stronger in females (Dearing and Gotlib, 2009; Hasselbalch et al., 2013; Joormann et al., 2007; see meta-analysis by Phillips et al., 2010). Due to the treatment-resistance (Bora et al., 2013; Hasselbalch et al., 2011) and the predictive nature of cognitive symptoms of MDD, researchers have called for further research on the cognitive symptoms of MDD (Prévot and Sibille, 2021; Romero et al., 2014).

Part of the challenge in MDD is the heterogeneity of symptoms. For example, sleep symptoms of MDD can be hypersomnia (~50% of episodes) or insomnia (~85% of episodes) and one third of individuals with MDD report instances of both (Geoffroy et al., 2018). The heterogeneity of symptoms is also coupled with sex differences in symptom severity, presentation, and comorbidities (Schuch et al., 2014). Not surprisingly, given the heterogeneitiy of MDD symptoms, there is heterogeneity in the underlying mechanisms of MDD (reviewed in Buch and Liston, 2021). Attention to this heterogeneity by examining the constellation of symptoms and further characterization of the mechanisms that drive these symptoms are needed across age and sex to improve the efficacy of future therapeutics (reviewed in Athira et al., 2020). Although there has been scant research in this area, there are profound sex differences in the neural and gene expression underpinnings following MDD (Gray et al., 2015; Labaka et al., 2018; reviewed in Bangasser and Cuarenta, 2021). These sex differences suggest that different treatment protocols may be needed to alleviate MDD in males versus females, but less research has examined sex by age differences.

Similar to humans, cognitive bias tasks are used in rodents to examine their affective state and negative cognitive bias is increased in rodent models of MDD and after exposure to aversive environments (Boleij et al., 2012; Burman et al., 2009, 2008; Enkel et al., 2010; Harding et al., 2004; Papciak et al., 2013; Richter et al., 2012). However, to our knowledge no studies have examined sex and age effects in response to cognitive bias testing. The neurobiology of negative cognitive bias is associated with structural and functional changes in the frontal cortex, hippocampus, amygdala, and nucleus accumbens in humans (Bijsterbosch et al., 2018; Browning et al., 2010; Poulsen et al., 2009; Sakaki et al., 2020; Siegle et al., 2007; Yang et al., 2010). Each of these brain regions are also disrupted in MDD (Kronenberg et al., 2009; McKinnon et al., 2009; Nauczyciel et al., 2013; Pizzagalli et al., 2009; van Eijndhoven et al., 2009), but more information is needed concerning the translational relevance of negative cognitive bias in rodents.

In the current study we sought to examine sex and age differences in cognitive bias and neural activation patterns across limbic, frontal, and reward regions. We developed a novel cognitive bias task linked to pattern separation, the ability to discriminate between highly similar situations (reviewed in Yassa and Stark, 2011). Pattern separation relies on hippocampal neurogenesis (Clelland et al., 2009; França et al., 2017; Sahay et al., 2011; Tronel et al., 2012) and both processes are impaired in MDD (reviewed in Gandy et al., 2017) and show sex differences (Yagi et al., 2016). We measured sex differences in cognitive bias and the expression of the immediate-early gene c-Fos, as a marker of neural activity (reviewed in Gallo et al., 2018; Kovács, 1998), in the subregions of the frontal cortex, hippocampus, amygdala, and nucleus accumbens in response to the display of cognitive bias in adolescent, young adult and middle-aged rodents. We hypothesized that negative cognitive bias and both neural activity and functional connectivity involved in the display of cognitive bias would differ by age and sex.

## 2. Methods and materials

### 2.1. Animals

Male and female Sprague-Dawley rats (N=95) were bred in house from animals obtained from Charles River (Québec, Canada). Only 1 male and 1 female rat per litter was assigned to each age group and each condition to avoid litter confounding effects. Males and females were housed (2-3 per cage) in separate colony rooms. Rats were maintained under a 12h light-dark cycle, with lights on at 07:00 h. Rats were housed in opaque polyurethane bins (48 × 27 × 20 cm) with aspen chip bedding and ad libitum access to autoclaved tap water and rat chow (Jamieson’s Pet Food Distributors Ltd, Delta, BC, Canada). Rats were left undisturbed, apart from weekly cage changing, until they reached the correct age for testing. All experimental procedures were approved by the University of British Columbia Animal Care Committee and in accordance with the Canadian Council on Animal Care guidelines.

### 2.2. Cognitive Bias Task Procedure

Male and female rats were randomly assigned to be tested in adolescence (postnatal day (PD) 40, *n*=29), young adulthood (PD 100, *n*=30), or middle-aged adulthood (PD 210, *n*=36) and then to one of the two groups - test rats (adolescents: male *n*=8, female *n*=9; young adults: *n*=9 per sex; middle-aged adults: *n*=12 per sex) and home cage controls (*n*=6 per sex per age).

Rats that underwent the 18-day cognitive bias procedure were placed in a shock-paired context (Context A) and in a no-shock-paired context (Context B) for 5 min each daily for 16 consecutive days, one context in the morning (8:30 h – 11:00 h) and the other context in the afternoon (13:00 h – 15:30 h). The placement of rats into Contexts A and B alternated between morning and afternoon each day of training (Fig. 1.). Test boxes used for all contexts had the dimensions of 30.5 × 24 × 21 cm (Med Associates Inc., St. Albans VT, USA) and they were cleaned with 70% isopropanol alcohol prior to testing each animal. Contexts A and B differed in several ways: illumination (one versus four lights illuminated), objects in the box (levers out or not), patterns on walls (distance between black lines on walls – 2mm or 15mm distance between lines), and transport time and method to the testing rooms (35 seconds or 1 min 35 seconds). In the shock-paired context, rats received three 0.6 milliamperes (mA) foot-shocks that lasted for 2 seconds with a 30-second inter-trial interval starting 3 min after being placed into the context box, with no-shock given in the other context. Whether Context A or Context B was paired with a footshock was counterbalanced between subjects. After 16 days of training, rats were placed in an ambiguous context (Context C) on Test Day (Day 18). Context C partially resembled both Contexts A and B in terms of transport (duration and method), illumination (two lights), one lever out, and an intermediate pattern of lines on the walls (7 mm between lines). Testing in Context C lasted 5 min with no footshock. Ninety min after exposure to Context C on day 18, test rats were euthanized by decapitation. Cage controls were left undisturbed except for weekly cage changing and were euthanized by decapitation age matched with the test rats. Brains were removed from the skull and cut in equal halves along the sagittal plane. The left hemisphere was used for c-Fos immunohistochemistry (described below).

**Fig. 1.**
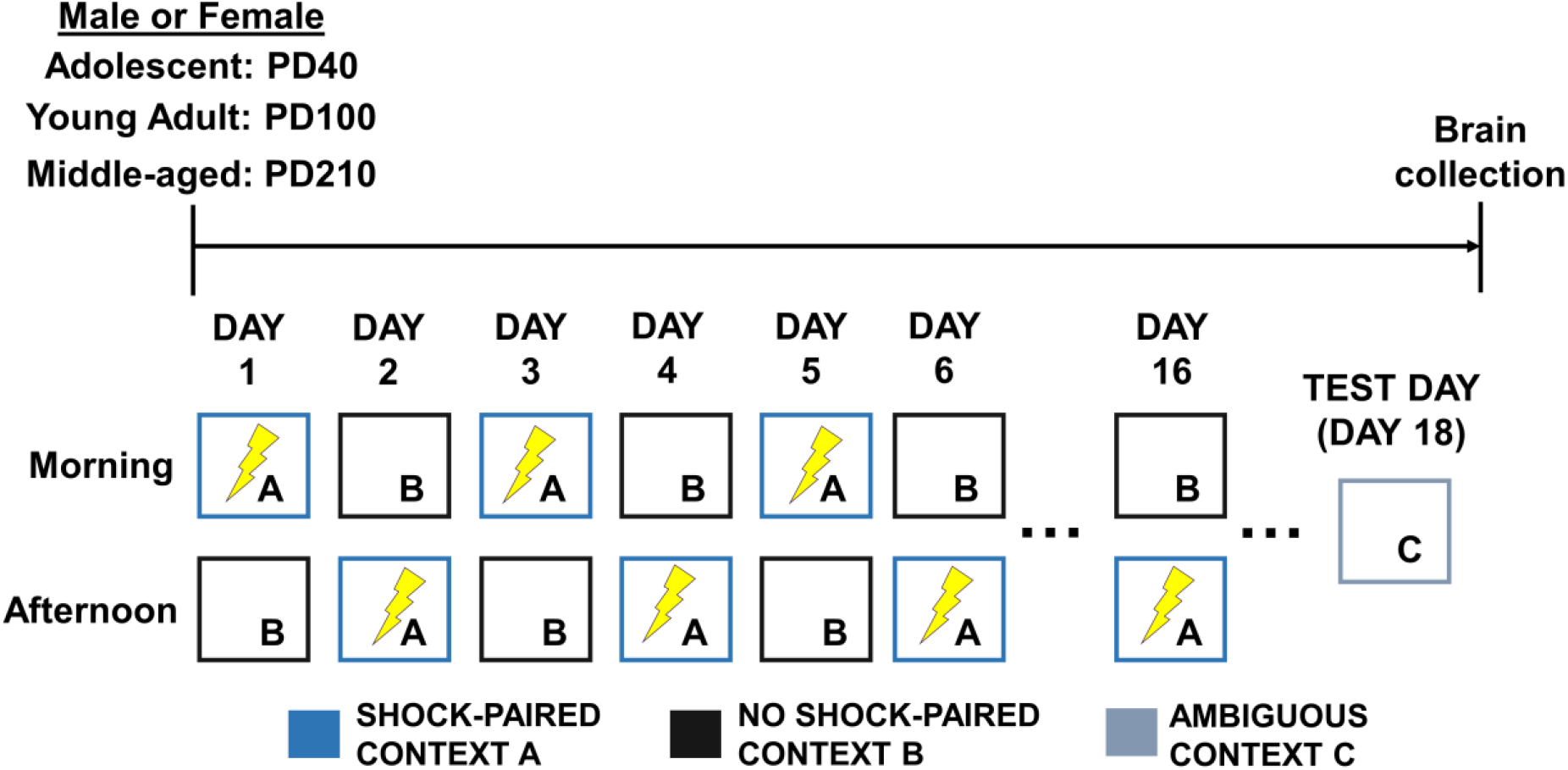
Experimental timeline and cognitive bias procedure.

### 2.3. Behavioral assessments

Rats were video recorded in the test chambers (Spy Store, Vancouver BC, Canada). Time spent freezing during the first 3 min of entering each context was measured on each day using the OpenFLD program (Johann, 2010) and percentage freezing was computed. Freezing behavior was defined as the rat displaying no head or body movement besides breathing (Barrientos et al., 2002). A difference score was created by subtracting percentage freezing in Context C on Day 18 from percentage freezing in Context B (no footshock-paired) on Day 16 and used to index negative cognitive bias scores (high freezing = negative cognitive bias; low freezing = neutral/positive cognitive bias; adapted from Hinchcliffe et al., 2017; Stuart et al., 2017).

A time sampling method was used to record behaviors every 10 seconds commencing 5 seconds after rats were placed into Context C, for a total of 19 observations per animal during the first 3 min of entering Context C (the ambiguous context). Behaviors examined included relaxed posture (no movement besides breathing), head movement (head moving side-to-side but no body movement), wall sniffing (sniffing the wall of the chamber), floor sniffing (sniffing the floor of the chamber), rotate body (in place rotating body around), rearing (rearing up, with weight on hind paws), allogroom (grooming self), and darting (a rapid forward movement across chamber). Behaviors were then divided into resting behavior (relaxed posture, head movement), active behavior (wall and floor sniffing, rotate body, rearing), and darting. The percentage of time that each rat engaged in each specific behavior was calculated.

### 2.4. c-Fos Immunohistochemistry

c-Fos is the protein of the immediate-early-gene c-fos that is transiently expressed in cells in response to action potentials and used as a marker of neuronal activation (reviewed in Kovács, 1998). The left hemisphere was placed into a 4% paraformaldehyde solution for 24 h, and subsequently placed into a 30% sucrose in 0.1M phosphate buffered saline (PBS; pH 7.4) for another 24 h and then until sliced. Coronal sections (30µm) were sliced on a microtome and collected from approximately bregma 3.72mm to −6.96mm (Paxinos and Watson, 2005). Sections were stored in an antifreeze solution (30% ethylene glycol, 20% glycerol in 0.1M phosphate buffer (PB; pH 7.4)) at −20 °C until immunohistochemistry assays were conducted.

Coronal sections were successively washed 4x in PBS for 10 min per wash and incubated at room temperature in a 0.6% hydrogen peroxide (H_2_O_2_; H1009, Sigma-Alrich, St. Louis, MO, USA) in distilled water (dH_2_O) for 30 min. Sections were then washed another 3x in 0.1M PBS for 10 min per wash, and then incubated at 4°C in c-Fos primary antibody (1:4000 Anti-c-Fos rabbit pAb; 190289; Abcam, Toronto, ON) for 24 hours. The next day, sections were washed 5x in 0.1M PBS for 10 min per wash and incubated overnight at 4°C in secondary antibody (biotinylated goat anti-rabbit IgG; 1:500; Vector Laboratories, Inc, Burlingame, CA). The last day, after another series of 5 washes in 0.1M PBS for 10 min per wash, sections were incubated in an avidin-biotin horseradish peroxidase solution (PK-4000, Vector Laboratories, Inc, Burlingame, CA) for 4 h at room temperature. Sections were washed 3x in 0.1M PBS for 10 min per wash and horseradish peroxidase was visualized using 3,3’ diaminobenzidine (DAB) in a 3 M sodium acetate buffer containing 2.5% nickel sulfate and 0.05% H_2_O_2_ (SK-4100, Vector Laboratories, Inc, Burlingame, CA) for 3 min. Sections were washed another 3x in 0.1M PBS for 10 min per wash and then mounted on Superfrost Plus slides (Fisher Scientific, Inc., Hampton, NH), let dry, dehydrated using increasing concentrations of ethanol (50%, 70%, 95%, 100% for 2, 2, 2, and 10 mins respectively), and then cleared with xylene for 10 min and coverslipped using Permount mounting medium (Fisher Scientific, Inc., Hampton, NH).

c-Fos protein immunostained brain sections were analyzed using a Nikon Eclipse 80i microscope. The retrieval of fear memories activates the hippocampus, frontal cortex, amygdala and nucleus accumbens (Blume et al., 2017; Gresack et al., 2009; Jin and Maren, 2015; Keiser et al., 2017; Maren et al., 1994; Piantadosi et al., 2020; Pothuizen et al., 2005; Quirk et al., 2003; Santarelli et al., 2018; Schmidt et al., 2019; Sotres-Bayon et al., 2012). Thus, digital images of regions of interest included the frontal cortex (anterior cingulate cortex = ACC, prelimbic cortex = PRL, infralimbic cortex = IL) within bregma 3.72 mm and 2.52 mm, the nucleus accumbens (nucleus accumbenc core = NAC, nucleus accumbens shell = NAS) within bregma 1.92 mm and 0.96 mm, the amygdala (basolateral amygdala = BLA, lateral amygdala = LA, central amygdala = CeA) within −2.16 mm and −2.92 mm, the dorsal hippocampus (dHPC; dentate gyrus (DG), CA3, CA1) within −2.64 mm and −4.56 mm, and the ventral hippocampus (vHPC; DG, CA3, CA1) within −5.76 mm and −6.36 mm were taken using 4x and 10x objectives. Photomicrographs were used to trace outline of each subregion of interest to calculate the area of each region using ImageJ software (Image J, 2020). Cell counts of c-Fos immunoreactive (ir) cells were conducted by experimenters’ blind to experimental condition and averaged across 4 sections per animal per subregion of interest using a 10x objective. c-Fos-ir cell density for each subregion of interest was calculated by dividing the cell count by the corresponding area in mm^2^ for each animal.

### 2.5. Data Analyses

Freezing was analyzed using a repeated measures analysis of variance (ANOVA) with context (shock, no-shock), day (1-16) as a within-subjects factors and with sex (male, female) and age (adolescence, young adulthood, middle-aged adulthood) as between-subjects factors. General linear mixed model ANOVAs with the same between-subjects factors above were performed on negative cognitive bias scores. Repeated-measures ANOVA on c-Fos density with subregion (frontal cortex: ACC, PRL, IL; nucleus accumbens: NAC, NAS; amygdala: BLA, LA, CeA; dorsal hippocampus: DG, CA3, CA1; ventral hippocampus: DG, CA3, CA1) and condition (homecage controls, test rats) as within-subjects factors with the between-subjects factors of age and sex were also conducted. Pearson product-moment correlations were conducted between c-Fos density in each region of interest and freezing in the ambiguous context. Principle component analyses (PCAs) were performed on the c-Fos data. Missing values, due to outliers or damaged tissue, which accounted for 7% of the data (2.2% outlier, 4.8% damaged tissue), were replaced by the mean for PCA analyses. Post-hoc tests used Newman-Keuls comparisons. Any *a priori* comparisons examining sex differences were subjected to Bonferroni correction. Significance level of p<0.05 was used. All statistical analyses were performed using Statistica software (v. 9, StatSoft, Inc., Tulsa, OK, USA).

Four animals were excluded from the following analyses due to their inability to distinguish between the shock- and no-shock-paired contexts on Day 16 of training (2 middle-aged males, 1 middle-aged female,1 young adult male).

## 3. Results

### 3.1. All animals, regardless of age and sex, learned to discriminate between the two contexts

All animals, regardless of age and sex, learned to discriminate between the shock-paired and no-shock-paired contexts (despite the four way interaction between day, context, sex, and age, F(30, 720) = 1.479, p = 0.049, *η*_*p*_^*2*^ = 0.058). Adolescent females and males discriminated between the two contexts on days 10-16 (all p’s<0.003 for females and p’s<0.017 for males with exception of day 13 which was p=0.07). Young adult females discriminated between the two contexts on from day 10-16 (p’s<0.0001; except days 11 and 13) and young adult males discriminated on days 12-16 (p’s<0.0001). In middle-age, females and males discriminated between the two contexts on days 12-16 (females: p’s<0.0002; males: p’s<0.019, except day 15). On day 1 of training, middle-aged males and females spent significantly more time freezing in the no-shock-paired context after exposure to the shock-paired context for the first time (p’s<0.0015) as well as young adult males (p=0.047). See Fig. 2.

**Fig. 2.**
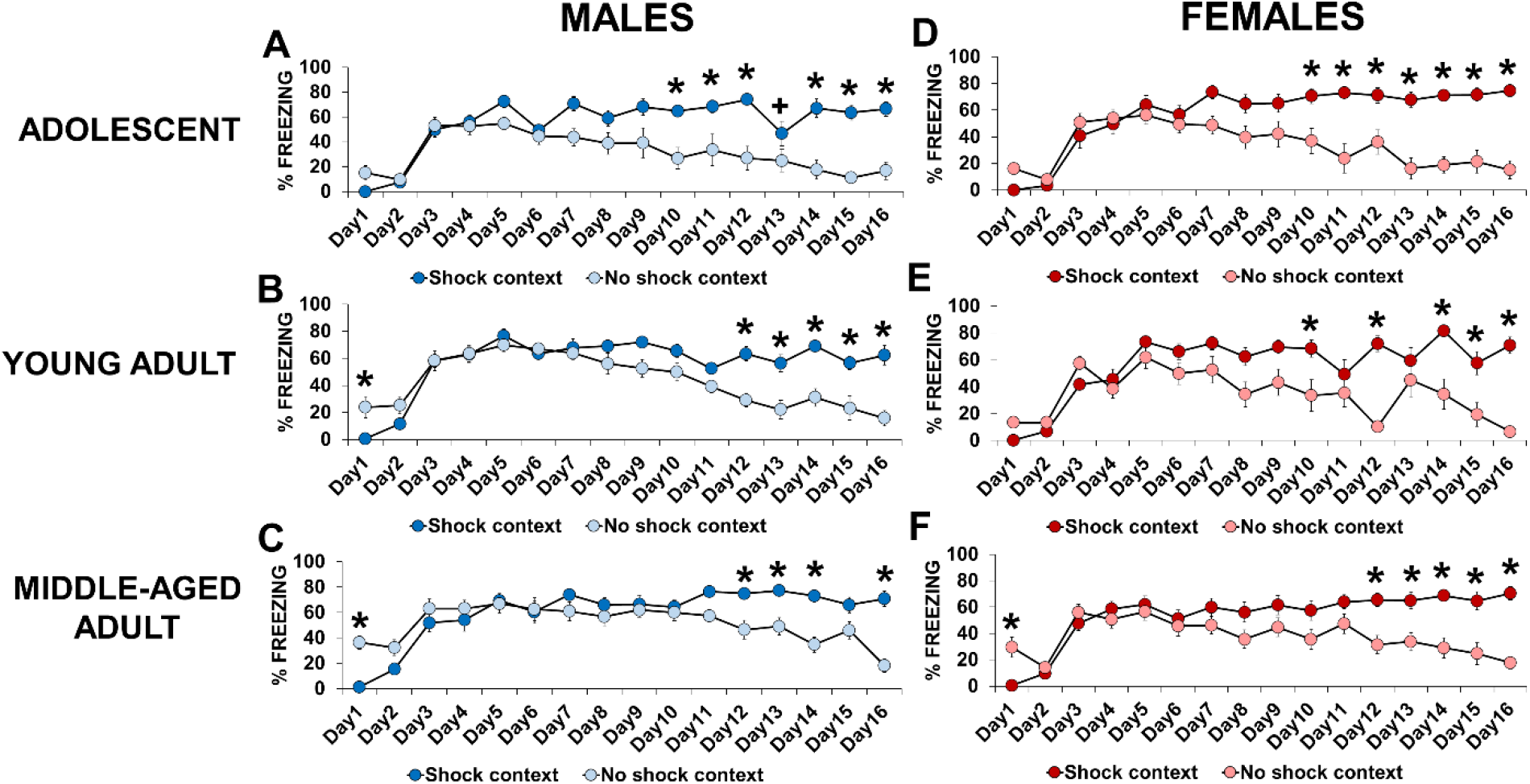
Percentage of time spent freezing in the shock and no-shock paired contexts during the 16 days of cognitive bias training in adolescent, young adult, and middle-aged males (**A-C**) and females (**D-F**). All rats learned to discriminate between the two contexts. On day 1, middle-aged males and females and young adult males spent more time freezing in the no-shock-paired context after experiencing the shock-paired context for the very first time. *indicates p<0.05 and +indicates p<0.1: difference between the contexts. *n*=7-11 per group.

### 3.2. Adults have a greater negative cognitive bias than adolescents, and males have a greater negative cognitive bias than females only in middle-age

We examined cognitive bias in two ways. First we compared freezing on day 16 in the shock-paired and no-shock-paired contexts to freezing in the ambiguous context on day 18 and then using a discrimination score. Freezing between the contexts differed depending on age (Context by age interaction: F(4, 98) = 2.527, p = 0.046, *η*_*p*_^*2*^ = 0.094). All rats spent more time freezing in the shock-paired context compared to the no-shock-paired (p’s<0.0002) and ambiguous contexts (p’s<0.04) regardless of sex and age. However, only middle-aged males and young adult females spent more time freezing in the ambiguous context compared to the no-shock-paired context (p=0.0003 and p=0.008, respectively). See Fig. 3A.

**Fig. 3.**
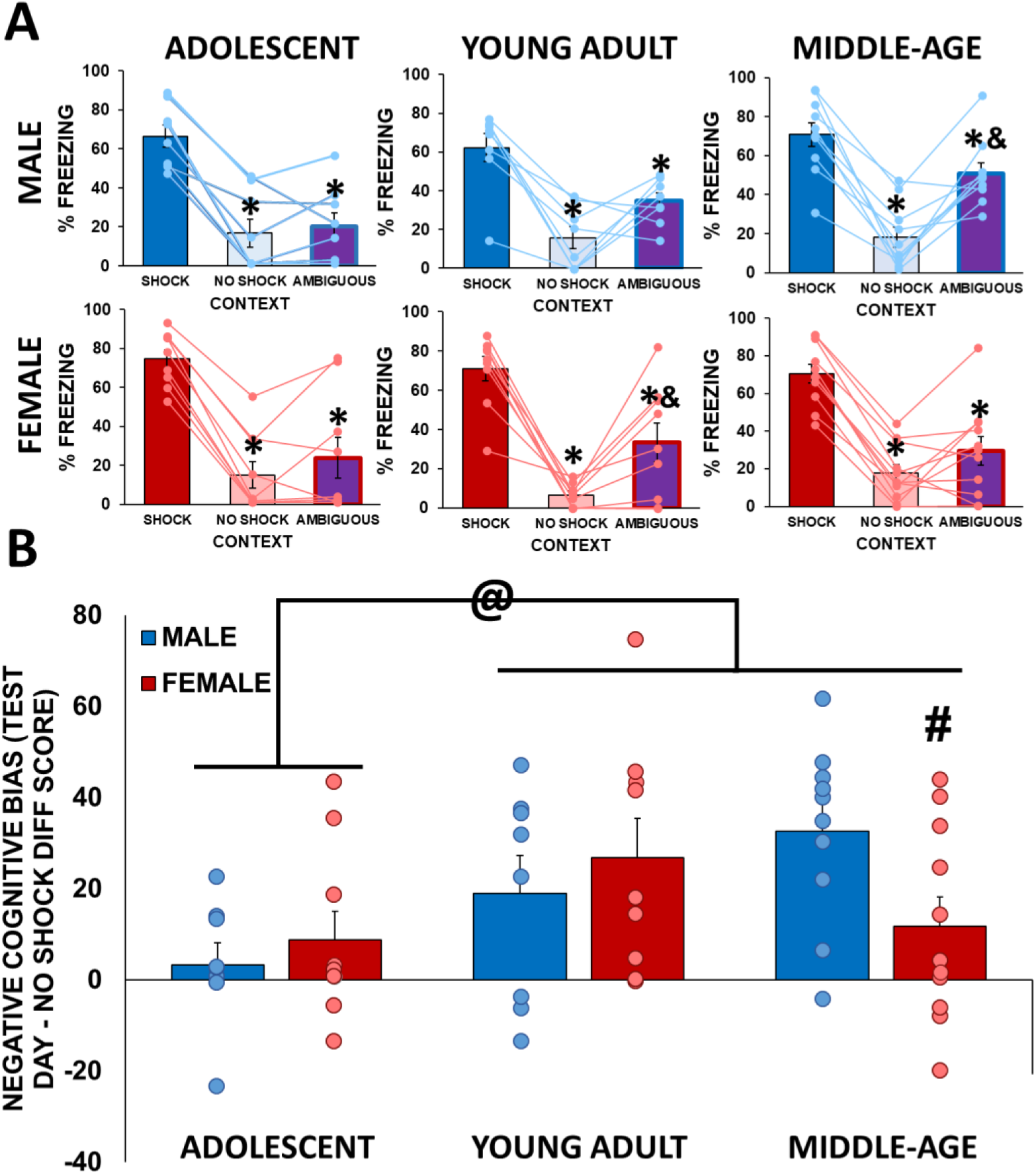
Percentage of time spent freezing in the shock and no-shock paired contexts on day 16 and in the ambiguous context on day 18 **(A)**. All groups spent more time freezing in the shock-paired context compared to the no-shock-paired and ambiguous contexts. Only young adult females and middle-aged males spent more time freezing in the ambiguous context compared to the no-shock-paired context. *indicates p<0.05: difference between the shock-paired context and other contexts. &indicates p<0.05: difference between ambiguous and no-shock-paired contexts. Negative cognitive bias discrimination scores (**B**). Young adults and middle-aged rats had greater negative cognitive bias scores than adolescents, and middle-aged males had greater negative cognitive bias than did middle-aged females. @indicates p=0.033: main effect of age. #indicates p=0.027: difference between middle-aged males and females. n=8-11 per group.

Both young adults and middle-aged adults had a greater negative bias than adolescents using the negative cognitive bias discrimination score (p’s <0.047; main effect of age: F(2, 49) = 3.64, p = 0.033, *η*_*p*_^*2*^ = 0.129). A priori we were interested in any sex differences, and in middle-age males displayed more negative bias than females (p = 0.027, cohen’s d = 1.014; sex by age interaction approached significance, F(2, 49) = 2.813, p = 0.070, *η*_*p*_^*2*^ = 0.103) which was not seen adolescents or young adults. See Fig. 3B.

### 3.3. Females displayed more active behavior than males

Females spent a higher percentage of time displaying active behavior compared to males (Main effect of sex: F(1, 49) = 4.254, p = 0.044, *η*_*p*_^*2*^ = 0.08). Males spent more time displaying resting behavior compared to females (Main effect of sex: F(1, 49) = 4.31, p = 0.043, *η*_*p*_^*2*^ = 0.08). Rats did not differ by age or sex in darting behavior (p’s>0.416). See Supplemental Fig. S1.

### 3.4. Females had greater neural activation than males and younger rats had greater neural activation to cognitive bias testing than older rats that depended on region

#### 3.4.1. Adult rats had higher c-Fos expression in the frontal cortex compared to adolescent rats

Young and middle-aged adult rats, regardless of sex, but not adolescent rats, had increased c-Fos expression in all frontal cortex subregions compared to controls (p’s < 0.00003; region by age by condition interaction: F(4,132)= 2.437, p=0.050, η_p_^2^ = 0.069). Young and middle-aged adult rats had higher c-Fos expression than adolescents in all frontal cortex subregions after exposure to the ambiguous context (p’s<0.007). See Fig. 4A-C.

**Fig. 4.**
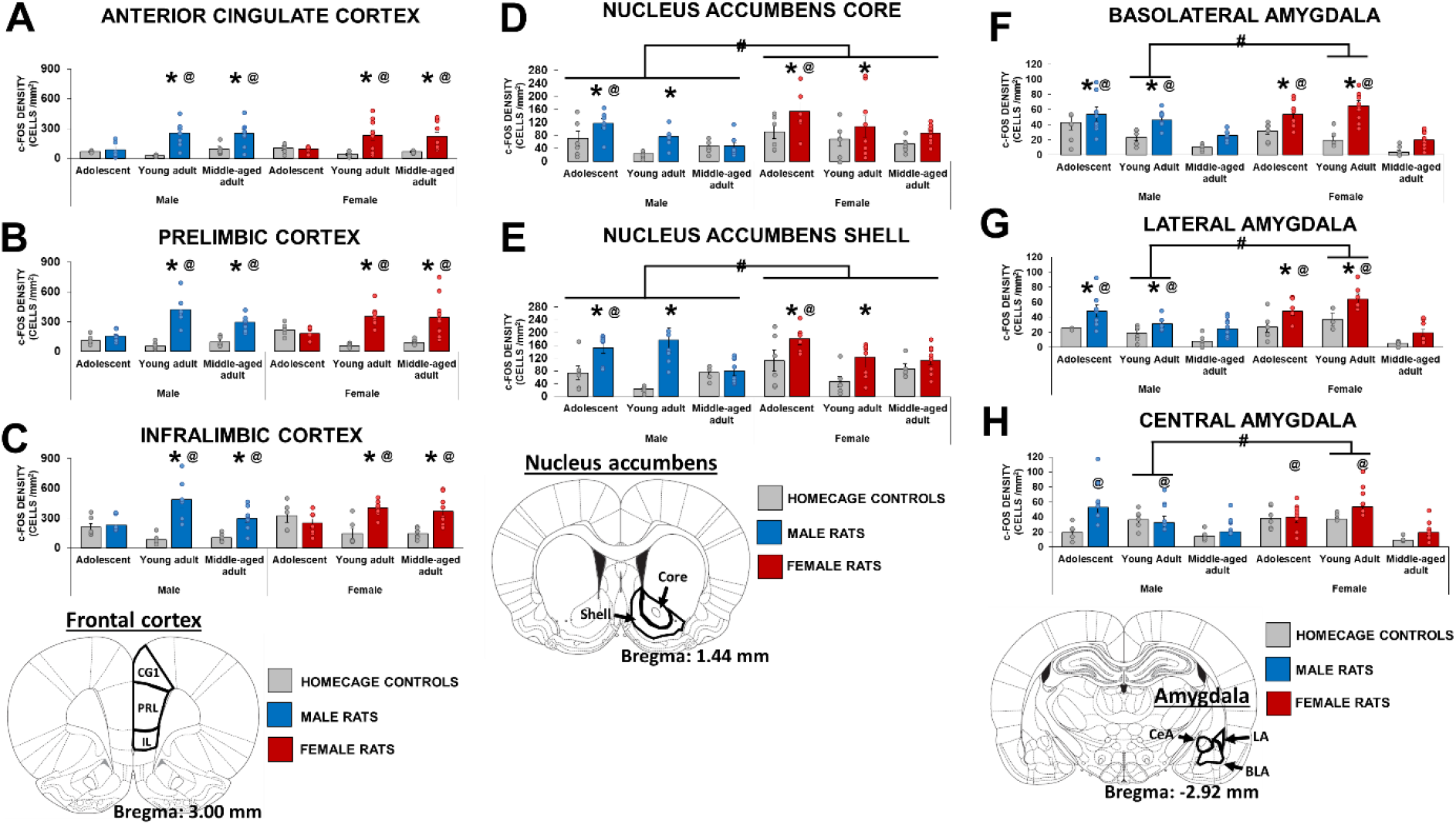
Young adults and middle-aged adults had higher c-Fos expression compared to adolescents and controls in the anterior cingulate cortex (**A**), prelimbic (**B**), and infralimbic (**C**) subregions of the frontal cortex. Females had higher c-Fos expression compared to males, adolescents had higher c-Fos expression compared to middle-aged adults, and adolescents and young adults had higher c-Fos expression than controls in the nucleus accumbens core (**D**) and shell (**E**). Young adult females had higher c-Fos expression compared to young adult males and young adults and middle-aged adults had higher c-Fos expression than adolescents in the basolateral (**F**), lateral (**G**), and central (**H**) subregions of the amygdala. * indicates p<0.05: compared to controls. #indicates p<0.05, sex effect. @indicated p<0.05 compared to adolescents or middle-aged adults. n=3-6 for homecage groups, n=6-10 for test rat groups.

#### 3.4.2. Females had higher c-Fos expression in the nucleus accumbens compared to males, adolescents had higher c-Fos expression compared to middle-aged rats

Overall, females had higher c-Fos expression in the nucleus accumbens core and shell compared to males (main effect of sex: F(1,67)=4.525, p=0.037, ηp2 = 0.063). Furthermore, c-Fos expression in nucleus accumbens subregions was dependent on condition and age (region by age by condition interaction: F(2,67)= 3.409, p=0.039, ηp2 = 0.092). In response to the ambiguous context, adolescent rats had higher c-Fos expression in the nucleus accumbens core and shell compared to young adult and middle-aged rats, regardless of sex (all p’s<0.005). Moreover, both adolescents and young adults had higher c-Fos expression in response to the ambiguous context compared to age-matched controls (p’s<0.0007). See Fig. 4D-E.

#### 3.4.3. Young adult females had higher c-Fos expression in the amygdala compared to young adult males, and adolescents and young adults had higher c-Fos expression than middle-aged adults

Young adult females had higher levels of c-Fos expression than young adult males (p=0.000008), but this sex difference was not seen in middle-age or adolescence (p’s>0.111; sex by age interaction: F(2,63)=4.115, p=0.021, η_p_^2^ = 0.116). All rats had higher c-Fos expression in the basolateral amygdala and lateral amygdala compared to controls (p’s<0.01), but only adolescents had higher c-Fos expression than controls in the central amygdala (p=0.004; region by age by condition interaction: F(4,126)=2.459, p=0.049, η_p_^2^ = 0.073). In addition, adolescents and young adults had higher c-Fos expression in all regions compared to middle-aged rats (p’s<0.00003). See Fig. 4F-H.

#### 3.4.4. Females had higher c-Fos expression in the dentate gyrus compared to males

Females had higher c-Fos expression in the dentate gyrus, regardless of region, compared to males overall (main effect of sex: F(1,72)=7.164, p=0.009, η_p_^2^ = 0.09). All rats, regardless of age, had higher c-Fos expression in the dentate gyrus of the dorsal hippocampus compared to the ventral hippocampus (p’s<0.0005; region by age interaction: F(2,72)=5.84, p=0.005, η_p_^2^ = 0.14). In addition, all groups, regardless of age, had higher c-Fos expression compared to controls in the dentate gyrus of the dorsal hippocampus (p=0.0001) but not the ventral hippocampus (p=0.587; region by condition interaction: F(1,72)=9.55, p=0.003, η_p_^2^ = 0.117). See Fig. 5A-B.

**Fig. 5.**
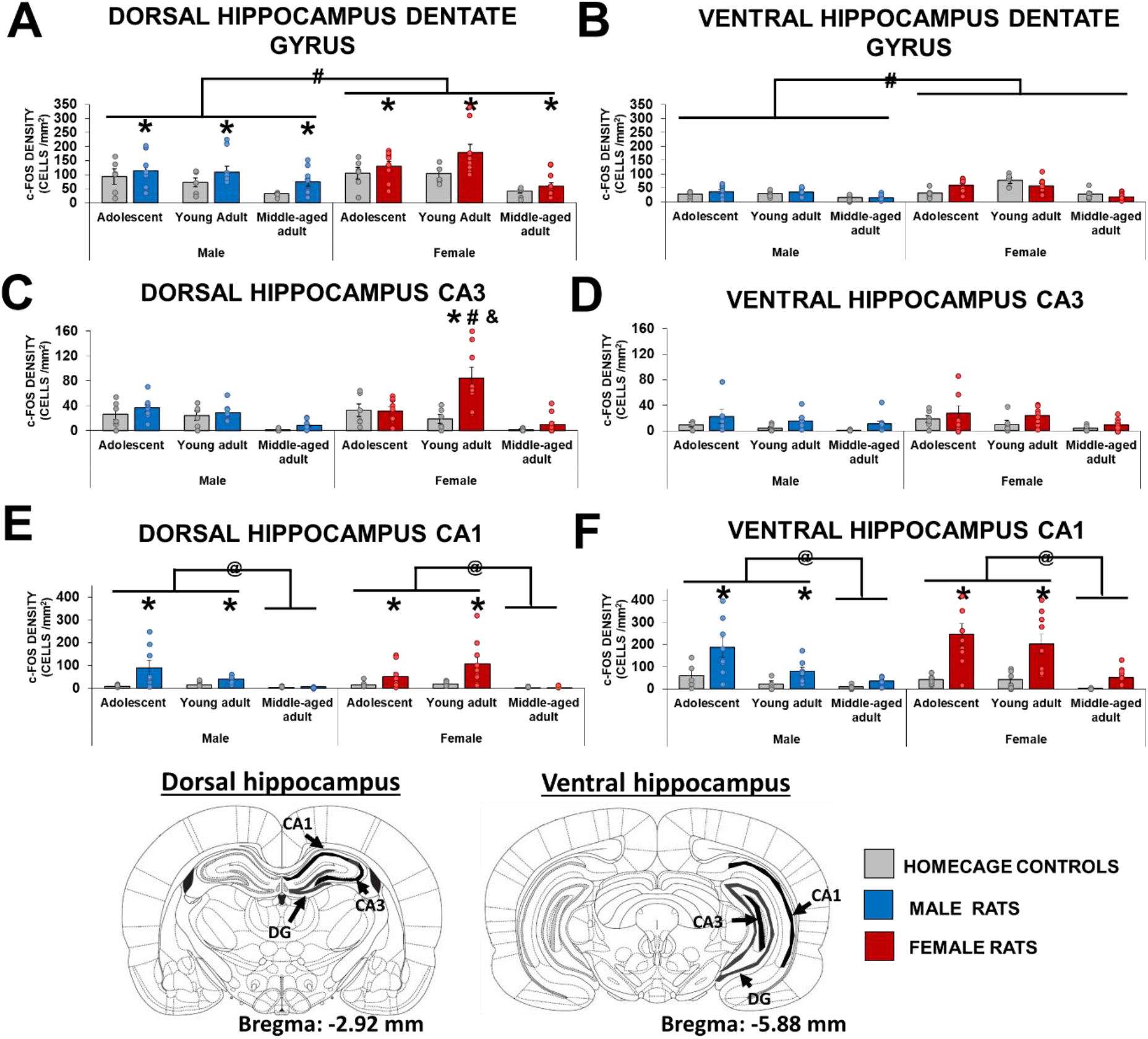
Mean (±SEM) density of c-Fos expressing cells in the dentate gyrus (A-B), CA3 (C-D), and CA1 (E-F) in the dorsal hippocampus and ventral hippocampus in male and female adolescents, young adults, and middle-aged adults. In the dentate gyrus, females had higher c-Fos expression compared to males overall, and all rats exposed to the ambiguous context had higher c-Fos expression in the dorsal dentate gyrus compared to controls. Young adult females had higher c-Fos expression in the dorsal CA3 compared to controls, young adult males, and c-Fos expression in the ventral CA3. Adolescents and young adults had higher c-Fos expression in the dorsal and ventral CA1 compared to middle-aged rats and controls after exposure to the ambiguous context. * indicates p<0.05: compared to controls. # indicates p<0.05: compared to the opposite sex. @ indicates p<0.05: compared to middle-age. & indicates compared to the ventral hippocampus. n=4-6 for controls, n=6-11 for test rats.

#### 3.4.5. Young adult females had higher c-Fos expression in the dorsal CA3 compared to young adult males

Young adult females had higher c-Fos expression in the dorsal CA3 region compared to young adult males (p=0.0001), controls (p=0.0002), and to c-Fos expression in the ventral CA3 region (p=0.0001) after exposure to the ambiguous context (region by sex by condition by age interaction: F(2,69)=3.971, p=0.023, η_p_^2^ = 0.103). See Fig. 5C-D.

#### 3.4.6. Adolescent and young adult rats had higher c-Fos expression in the CA1 region compared to controls and middle-aged rats

Adolescent and young adults, but not middle-aged rats, had higher c-Fos expression in the CA1 region, regardless of sex and dorsoventral axis compared to controls (p’s<0.0002; condition by age interaction: F(2,70)=5.687, p=0.005, η_p_^2^ = 0.14). Furthermore, adolescents and young adults had higher c-Fos expression compared to middle-aged rats in the CA1 region (p’s<0.0006). See Fig. 5E-F.

### 3.5. Freezing was negatively associated with c-Fos expression in females, but not in males, in the hippocampus and infralimbic cortex, dependent on age

Overall, there were more significant correlations with freezing in the ambiguous context and neural activation in females (6) compared to males (1), and the general pattern was negative correlations in females in the hippocampus and infralimbic cortex. High freezing behavior in response to the ambiguous context was associated with lower c-Fos expression in the hippocampus of adolescent females (dorsal CA3 (r = −0.762, p = 0.017) and in the hippocampus (dorsal DG (r = −0.668, p = 0.025) and ventral DG (r = −0.631, p = 0.037), ventral CA1 (r = −0.770, p = 0.006)) and infralimbic cortex (r = −0.652, p = 0.041) of middle-age females. However, there were two positive correlations as high freezing behavior in response to the ambiguous context was associated with higher c-Fos expression in the nucleus accumbens core of young adult males (r = 0.801, p = 0.05) and in the infralimbic cortex of young adult females (r = 0.705, p = 0.034). See Supplemental Fig. S2.

### 3.6. Functional connectivity in response to the ambiguous context depends on sex and age

We examined the effects of functional connectivity in two ways, using correlations of c-Fos expression between different brain regions and using a PCA to reduce the data set while preserving information.

When only examining significant correlations ≥ 0.7 (adapted from Wheeler et al., 2013) we found that hubs of correlations between activated regions shifted across age and sex as can be seen in Fig. 6. If we define a hub as 2 or more of these correlations between regions we see that both the hub location and the valence of correlations between activation levels across the multiple bran regions shifted with age and sex, suggesting that both of these are important factors in neural activation patterns in response to the ambiguous context.

**Fig. 6.**
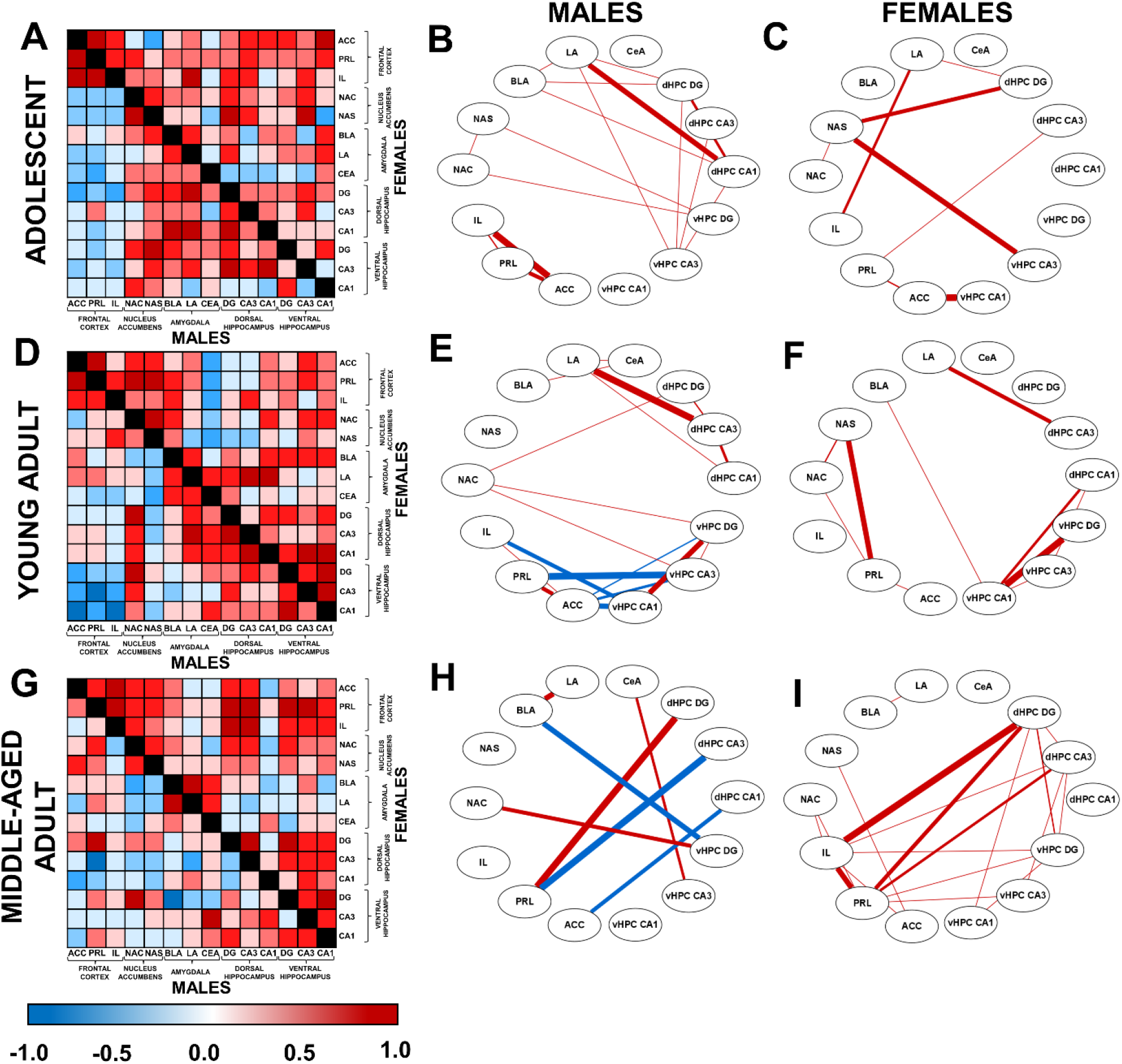
Heatmaps showing all correlations and cytoscape graphs showing significant correlations ≥ 0.7 between c-Fos expression in each region of interest in male and female adolescent (A-C), young adult (D-F), and middle-aged (G-I) rats in response to the ambiguous context. The thickness of the lines in B,C,E,F,H, and I are related to the strength of the correlation (stronger is thicker), whereas the color relates to the valence (positive (red) or negative (blue)) of the correlation. Only adult males (young adult, middle-age) had negative correlations of activation ≥ 0.7 between regions of the hippocampus and the frontal cortex or the BLA. In females, during adolescence the major hub was between the NAS and the hippocampus which shifted to a hub between the NAS and the frontal cortex in young and middle-aged adults. Note that between males and females there were distinct patterns of functional connectivity within the same age groups. Furthermore, several hubs between the frontal cortex and the hippocampus emerged in middle-age in males and females. Several correlations of activation ≥ 0.7 between the frontal cortex and the hippocampus emerged in middle-age. ACC = anterior cingulate cortex, PRL = prelimbic cortex, IL = infralimbic cortex, NAC = nucleus accumbens core, NAS = nucleus accumbens shell, BLA = basolateral amygdala, LA = lateral amygdala, CeA = central amygdala, DG = dentate gyrus, CA3 = cornu ammonis 3, CA1 = cornu ammonis 1.

There were significant sex differences in these correlations between activation in the frontal cortex and hippocampus in adolescence (ACC and ventral CA1 (z=1.754, p=0.04), IL and dorsal DG (; z=1.739, p=0.041)), young adulthood (ACC and ventral CA1 and CA3, p=0.027 and p=0.006 respectively; PRL and ventral CA1 and CA3, p=0.034 and p=0.006 respectively; IL and ventral CA1, z=-1.768, p=0.039) and in middle-age (PRL and dorsal CA3 (z=-4.398, p<0.0001; IL and dorsal DG (p<0.0001) and CA3 (p=0.016)).

### 3.7. Functional Connectivity between the limbic and reward regions during cognitive bias was influenced by sex and age

The first two principal components accounted for 59.7% of the variance of the c-Fos expression data. Component 1 accounts for 41.44% of the variance and has strong associations between the nucleus accumbens, amygdala, and hippocampus. Component 2 accounts for 18.26% of the variance and is associated with functional connectivity within the frontal cortex. Factor loadings for the principal components were subjected to an ANOVA and are shown in Table 1. An ANOVA on Principle Component 1 found that functional connectivity between all brain regions except the frontal cortex in response to cognitive bias was greater in young adult females compared to young adult males (p=0.0009) and greater in younger rats (adolescents, young adults) compared to middle-aged rats (p’s<0.022; age and sex interaction: F(2,48)=4.68, p=0.014, ηp2 = 0.163). An ANOVA on Principle Component 2 found that frontal cortex functional connectivity was greater in response to cognitive bias in adult rats (young adult, middle-aged) compared to adolescents (p’s<0.006; main effect of age: F(2,48)=11.55, p=0.00008, ηp2 = 0.325). See Fig. 7.

**Table 1.**
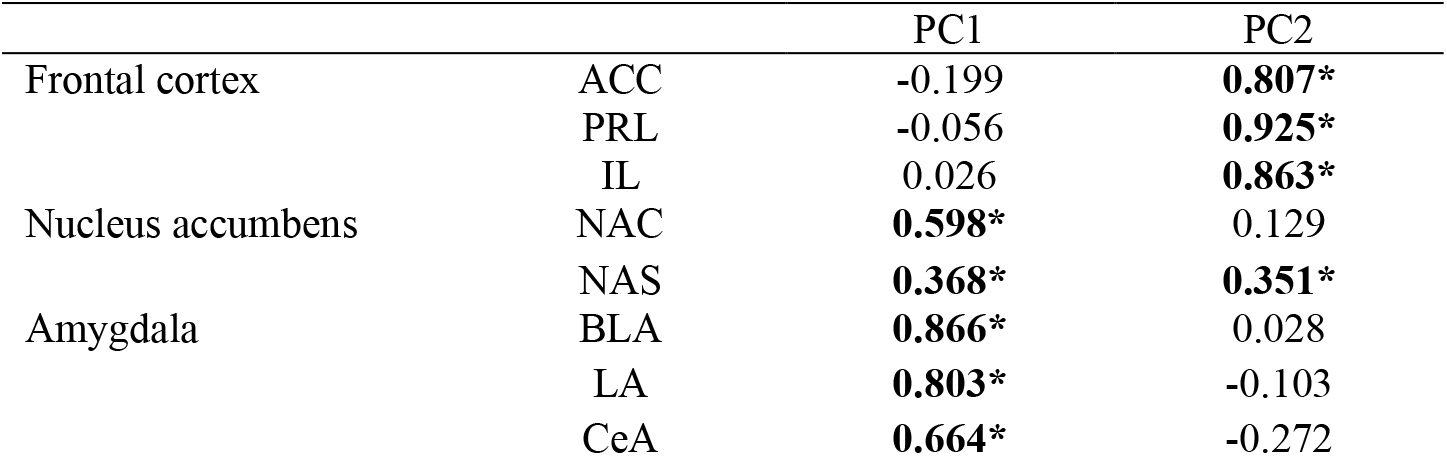
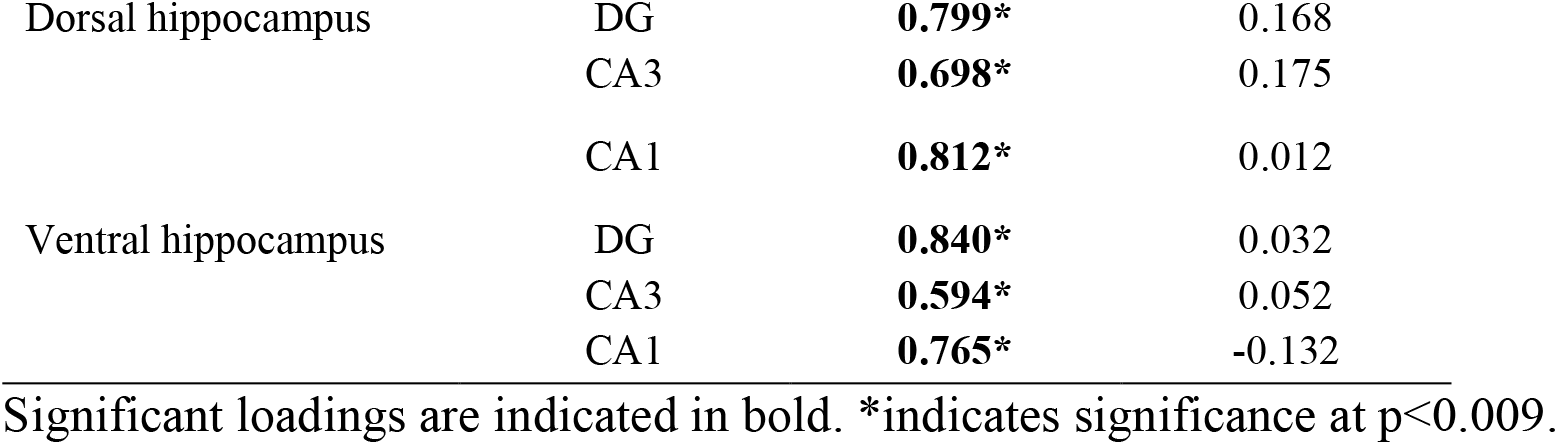
Principal component loadings from c-Fos expression

|  |  | PC1 | PC2 |
| --- | --- | --- | --- |
| Frontal cortex | ACC | -0.199 | **0.807\*** |
|  | PRL | -0.056 | **0.925\*** |
|  | IL | 0.026 | **0.863\*** |
| Nucleus accumbens | NAC | **0.598\*** | 0.129 |
|  | NAS | **0.368\*** | **0.351\*** |
| Amygdala | BLA | **0.866\*** | 0.028 |
|  | LA | **0.803\*** | -0.103 |
|  | CeA | **0.664\*** | -0.272 |
| Dorsal hippocampus | DG | **0.799\*** | 0.168 |
|  | CA3 | **0.698\*** | 0.175 |
|  | CA1 | **0.812\*** | 0.012 |
| Ventral hippocampus | DG | **0.840\*** | 0.032 |
|  | CA3 | **0.594\*** | 0.052 |
|  | CA1 | **0.765\*** | -0.132 |

Significant loadings are indicated in bold. \*indicates significance at $p < 0.009$ .

**Fig. 7.**
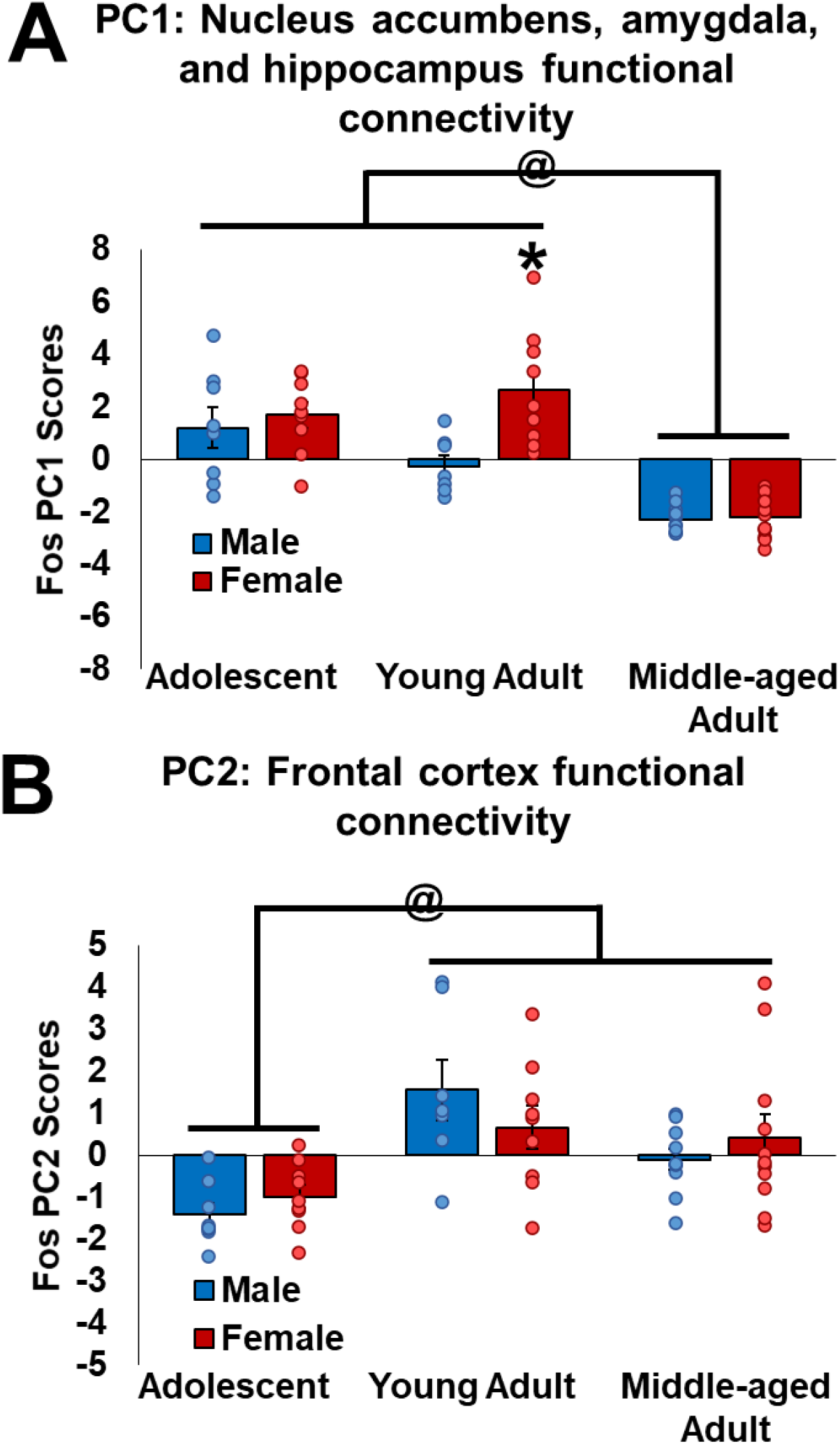
Principle component (PC) scores for c-Fos expression data. PC1 scores (**A**), associated with functional connectivity between the amygdala, nucleus accumbens, and hippocampus, was more involved with the cognitive bias of young adult females compared to young adult males, and in adolescents and young adults compared to middle-aged rats. PC2 scores (**B**), associated with frontal cortex functional connectivity, was more involved in the cognitive bias of adolescents and young adults. * indicates p<0.05: compared to young adult males. @indicates p<0.05: comparison between age groups. *n*=7-11per sex/age.

## 4. Discussion

Using our novel cognitive bias testing procedure, we report sex and age differences in cognitive bias behavior as well as neural activation in response to an ambiguous context. Adults (young adults and middle-aged) display greater negative cognitive bias compared to adolescents. A sex difference in behavior only emerged in middle-age, as males display greater negative cognitive bias than females. Despite the lack of sex differences in cognitive bias in adolescence and young adulthood, there were profound sex differences in neural activation following cognitive bias testing. Females had greater neural activation in the limbic and reward regions than males in response to cognitive bias testing. Age also played a role as, perhaps predictably, young rats had greater neural activation in response to cognitive bias testing in most regions than older rats. Interestingly, negative correlations between freezing behavior and neural activation in the hippocampus or frontal cortex were seen in females, suggesting greater activity was associated with a more positive/neutral bias in females, but not in males. Moreover, functional connectivity in response to the ambiguous context shifted with sex and age. All in all, our findings lay the groundwork for using cognitive bias testing to examine negative bias as a depressive-like endophenotype and affirms that sex and age are crucial factors to consider.

### 4.1. Negative cognitive bias increases with age, and middle-aged males have a greater negative cognitive bias than middle-aged females

Our results demonstrate that negative cognitive bias increased with age as young adult and middle-aged rats had a greater negative cognitive bias than adolescent rats. Our results are consistent with findings that adolescents are more optimistic in the literature (Rodham et al., 2006). Low freezing behavior in adolescents exposed to the ambiguous context is also similar to past findings of less freezing in adolescent male and female rats in novel environments compared to adults (Bronstein, 1972; Philpot and Wecker, 2008). Our results are not reflective of any difference in context discrimination as the adolescents learned to discriminate between the two contexts at same time point as adults. Collectively, these data suggest that a neutral/positive cognitive bias in adolescents may be attributed to immature risk assessment.

A sex difference emerged in with age as middle-aged males had a greater negative cognitive bias compared to middle-aged females. The linear increase in negative cognitive bias in males resembles findings of increased fixation on negative stimuli from young to old age in the human literature (sex not examined; Bucher et al., 2020). Our finding is also partially consistent with a study that used cued-fear fear conditioning, as male rats exhibited increased freezing in a novel context from adolescence to adulthood, an effect that was not seen in females (Colon et al., 2018). Possible explanations for the sex difference in middle-age are sex differences in fear behavior (reviewed in Tronson and Keiser, 2019), fear generalization, or in memory retrieval (Keiser et al., 2017). In terms of fear behavior, darting, an escape-like behavior, is an active fear response during cued-fear conditioning seen more in female compared to male rats (Colom-Lapetina et al., 2019; Gruene et al., 2015), which can explain low levels of freezing in females. However, we did not see any sex differences in darting in our data, consistent with observations that darting is observed in response to cued and not context fear conditioning (reviewed in Odynocki and Poulos, 2019). In terms of fear generalisation, if this was present in one sex over the other one might expect greater freezing to the non-shock context, but there were no sex differences in the amount of freezing in the non-shock context across any age group. Furthermore, regardless of age and sex, rats did not differ in their discrimination between the shock-paired and no-shock-paired contexts. These data suggest that differences in cognitive bias after exposure to the ambiguous context are not due to differences in contextual discrimination across age and sex. Thus, sex differences in fear expression or memory retrieval do not appear to be underlying the sex difference in negative cognitive bias in middle-age. Further studies should explore whether sex differences in negative cognitive bias in middle-aged rats remain when using other cognitive bias tasks such as the go/no-go or go/go tasks (reviewed in Nguyen, Guo, and Homberg, 2020).

### 4.2. Greater role of the nucleus accumbens, amygdala, and dentate gyrus in the cognitive bias of females compared to males

Females had higher c-Fos expression in the dorsal and ventral DG, nucleus accumbens, and in the amygdala compared to males, which, at least with respect to the amygdala, depended on age. Sex differences in mechanisms underlying fear memories have been observed in the hippocampus, amygdala, and the frontal cortex (Blume et al., 2017; Gresack et al., 2009; Keiser et al., 2017; Maren et al., 1994). Our data resemble past findings in mice of higher c-Fos expression in females than in males in the amygdala during fear retrieval but are in contrast to higher c-Fos expression in the dorsal hippocampus of males than females (Colon and Poulos, 2020; Keiser et al., 2017). However, another study, found that inactivation of adult-born neurons in the dentate gyrus decreased fear expression and enhanced contextual discrimination in adult female mice (Huckleberry et al., 2018). Greater amygdala activity was also associated with an increased negative cognitive bias in middle-aged postmenopausal human females with remitted MDD compared to females without a history of MDD (Albert et al., 2017), suggesting similar patterns may be seen in humans. Together, these studies suggest a major role of amygdala activation in female fear retrieval and negative cognitive bias and hippocampal activation in female fear expression and contextual discrimination.

### 4.3. The neural expression of cognitive bias shifts to involve the frontal cortex with age

Our finding of increased activation in the frontal cortex subregions of young adults and middle-aged adults but not adolescents suggest a greater involvement of these regions in the cognitive bias of adults. Indeed, increased IEG expression and functional connectivity between subregions of the frontal cortex (ACC, PRL, IL) are involved in distinguishing between fear-related and novel contexts in adult male mice and rats (Chakraborty et al., 2016; Frankland et al., 2004; Zelikowsky et al., 2014, 2013). In adolescent rodents, there is less of a role of PRL and IL activity in contextual fear (Heroux et al., 2017; but see Samifanni et al., 2021) and this may be linked to a reduced modulation of the amygdala by the PRL and IL in adolescence compared to adulthood (Gee et al., 2013; Selleck et al., 2018). Thus, reduced neural activity in the subregions of the frontal cortex of adolescents compared to adults may interact with contextual fear to modulate age differences in the display of cognitive bias.

### 4.4. Negative correlations between freezing and hippocampal activity in adolescent and middle-aged females

Intriguingly in our present data we saw negative correlations between freezing in the ambiguous context and neural activation in adolescent and middle-aged females, suggesting that negative cognitive bias was related to less activity in the IL, ventral DG and CA1, and dorsal CA3. Our results are partially consistent with findings of sex differences in the involvement of the ventral hippocampus in contextual fear, as activity in the ventral hippocampus is positively correlated with contextual fear in males, but not females (Gresack et al., 2009). In addition, we found that correlation hubs between activated regions in response to the ambiguous context shifted depending on age and sex. The number of hubs that included correlations between activation in the frontal cortex and limbic regions increased in adulthood and adult males had negative correlation hubs between activation in the frontal cortex and the hippocampus that were not seen in females. These data suggest age and sex differences in how neural activity is involved in cognitive bias and underscore that treatments influencing neural communication may need to be tailored to sex and age.

### 4.5. Importance of Cognitive Bias as a Symptom of MDD

MDD is a complex condition and cognitive systems involved in MDD have been overlooked. In contrast with other cognitive bias procedures in rodents (Brydges et al., 2012, 2011; Brydges and Hall, 2017; Chaby et al., 2013; Hinchcliffe et al., 2017; Jones et al., 2018; Stuart et al., 2017), our procedure is not based on active choice/motivation and measures the degree of negative cognitive bias by targeting the evaluation of an ambiguous situation as either negative or neutral. Furthermore, we do not see any age or sex differences in the rats discriminating between the two contexts during the training phase which gives us confidence that the age by sex changes we see in negative cognitive bias during the test phase are not due to mnemonic differences. Our findings parallel the human literature that show the involvement of the frontal cortex, amygdala, hippocampus, and nucleus accumbens in cognitive bias (Hilland et al., 2020; Sakaki et al., 2020) (Sakaki et al., 2020; Hilland et al., 2020; reviewed in Mineur and Picciotto, 2019) showing translational and construct relevance. We also found that our cognitive bias procedure is less time consuming as rats quickly (within 16 days) achieve stable levels of performance, across the lifespan, compared to other cognitive bias tasks (which take several weeks; reviewed in Bethell, 2015). Future studies will examine negative cognitive bias in rodent models of MDD and the molecular underpinnings of negative cognitive bias between males and females across age.

## 5. Conclusions

Overall, we found sex and age differences in rat cognitive bias using a novel paradigm and in functional connectivity involved in this bias, despite similar ability to discriminate between the two contexts. Negative cognitive bias increased from adolescence to adulthood and middle-aged males had a greater negative cognitive bias than middle-aged females. The nucleus accumbens, amygdala, and hippocampal regions showed greater involvement in the cognitive bias of females than males and in the younger ages compared to the middle-aged rats. The frontal cortex showed greater involvement in the cognitive bias of adults compared to adolescents. Given that cognitive dysfunction in MDD has been a target of therapy and negative cognitive bias remains treatment resistant we hope our novel cognitive bias procedure will be useful for examining novel therapeutics for MDD and negative cognitive bias in animal models of depression. Our findings also stress the importance of considering sex and age when examining depressive symptoms. In addition, these data provide the groundwork needed to determine what brain regions and patterns of functional connectivity between regions should be targeted when investigating sex and age-specific cognitive bias. Understanding the underlying neural mechanisms of negative cognitive bias will bring us closer to precision medicine for treatment-resistant negative cognitive bias.

## Supporting information

Supplemental Section

## Author contribution statement

**Travis Hodges**: Conceptualization, Methodology, Formal analysis, Writing – Original draft, Review, & Editing, Investigation. **Liisa Galea:** Conceptualization, Methodology, Formal analysis, Writing – Original draft, Review, & Editing, Supervision. **Grace Lee:** Investigation. **Sophia Noh**: Investigation.

## Acknowledgments

Financial support for this research was provided by an operating grant from the Canadian Institutes for Health Research (MOP142328) to LAMG. TEH was supported by the University of British Columbia Institute of Mental Health Marshalls Scholars Program.

