## Supplemental Section for "Sex and age differences in cognitive bias and neural activation in response to cognitive bias"

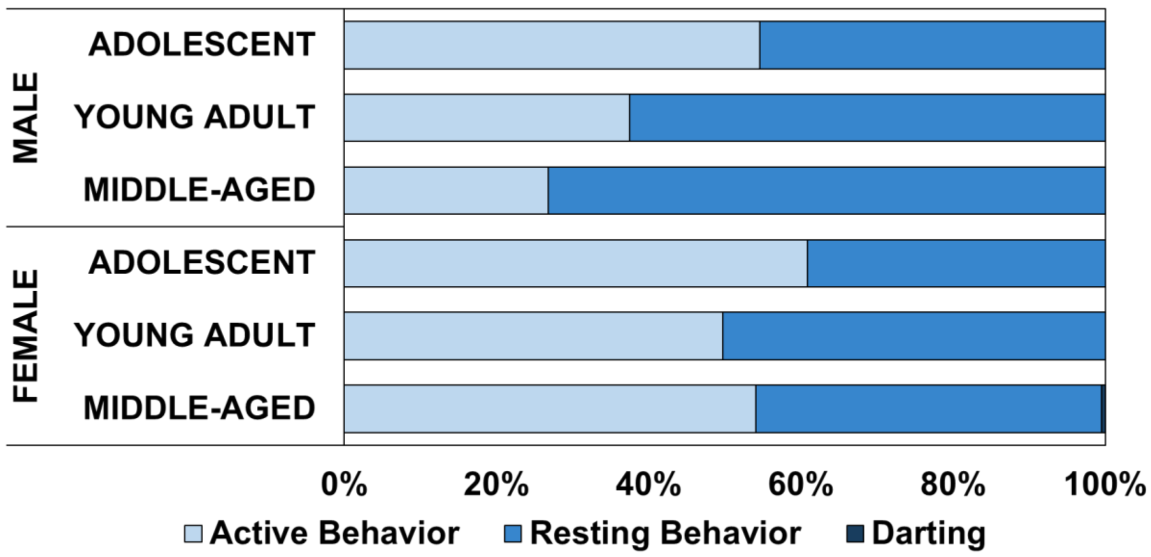


**Fig. S1.** Percentage of time spent displaying active behavior, resting behavior, or darting in the ambiguous context. Females spent more time displaying active behaviors whereas males spent more time displaying resting behaviors. Only middle-aged females spent 0.48% of time displaying darting behavior in the ambiguous context.

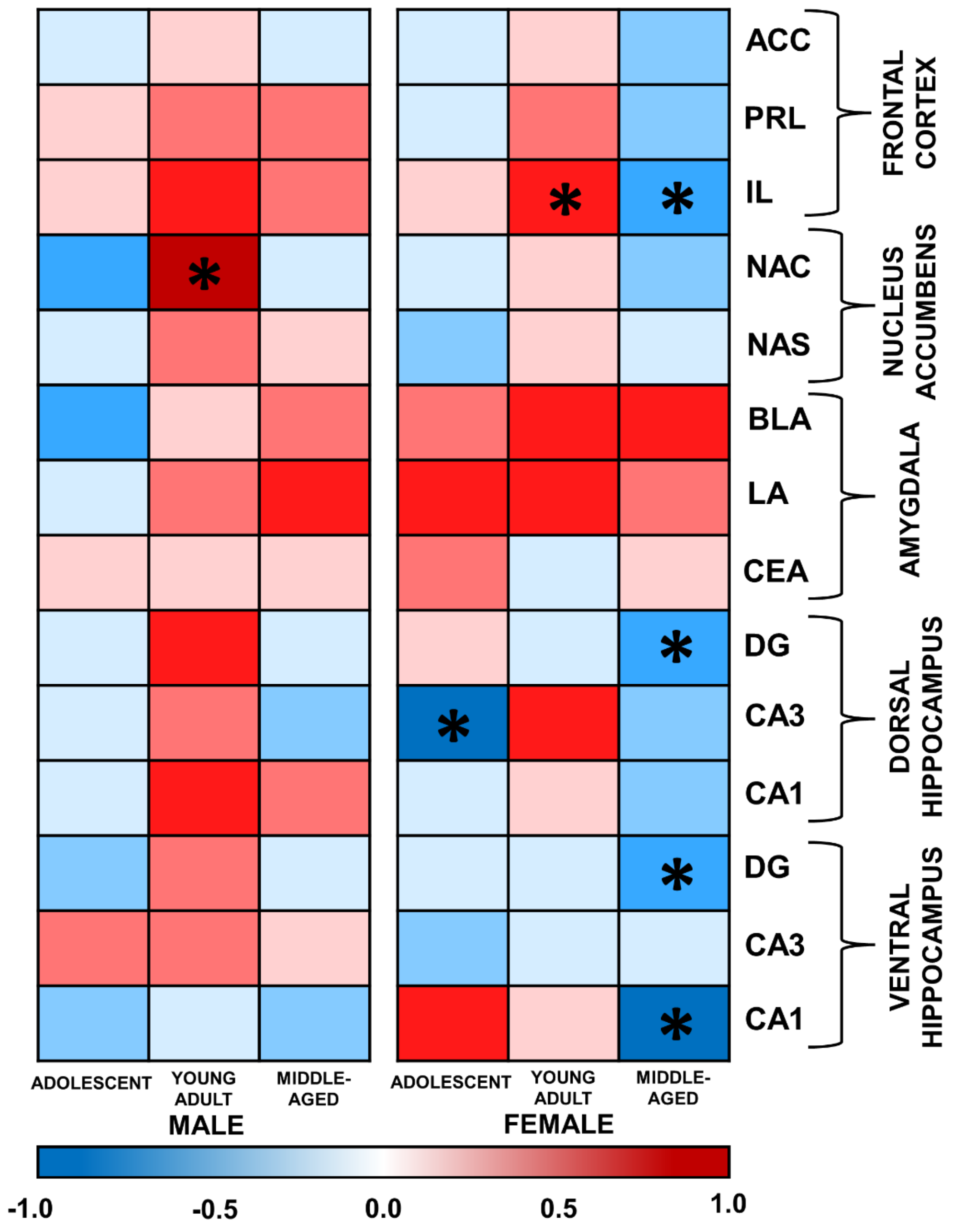


**Fig. S2.** Heatmaps showing correlations between neural activity in each region and freezing behavior in the ambiguous context. High freezing behavior in response to the ambiguous context was associated with lower c-Fos expression in the hippocampus of adolescent females and in the hippocampus and IL of middle-age females. High freezing behavior in response to the ambiguous context was associated with c-Fos expression in the NAC of young adult males and in the IL of young adult females. * indicates significant correlation $p<0.05$.
